# Patulin contamination of hard apple cider by Paecilomyces niveus and other postharvest apple pathogens: assessing risk factors

**DOI:** 10.1101/2023.01.16.523797

**Authors:** Tristan W. Wang, Amanda G. Wilson, Gregory M. Peck, Patrick A. Gibney, Kathie T. Hodge

## Abstract

Hard apple cider is considered to be a low-risk product for food spoilage and mycotoxin contamination due to its alcoholic nature and associated food sanitation measures. However, the thermotolerant mycotoxin-producing fungus *Paecilomyces niveus* may pose a significant threat to hard cider producers. *Pa. niveus* is known to infect apples (*Malus* x*domestica*), and previous research indicates that it can survive thermal processing and contaminate finished apple juice with the mycotoxin patulin. To determine if hard apple cider is susceptible to a similar spoilage phenomenon, cider apples were infected with *Pa. niveus* or one of three patulin-producing *Penicillium* species and the infected fruits underwent benchtop fermentation. Cider was made with lab inoculated Dabinett and Medaille d’Or apple cultivars, and patulin was quantified before and after fermentation. Results show that all four fungi can infect cider apples and produce patulin, some of which is lost during fermentation. Only *Pa. niveus* was able to actively grow throughout the fermentation process. To determine if apple cider can be treated to hinder *Pa. niveus* growth, selected industry-grade sanitation measures were tested, including chemical preservatives and pasteurization. High concentrations of preservatives inhibited *Pa. niveus* growth, but apple cider flash pasteurization was not found to significantly impact spore germination. This study confirms that hard apple cider is susceptible to fungal-mediated spoilage and patulin contamination. *Pa. niveus* should be of great concern to hard apple cider producers due to its demonstrated thermotolerance, survival in fermentative environments, and resistance to sanitation measures.

**Highlights:**

- Apple fruits of traditional cider cultivars Dabinett and Medaille d’Or were found to be susceptible to infection by three patulin-producing *Penicillium* spp. and *Paecilomyces niveus*
- *Pa. niveus* can grow in finished fermented hard cider at 5.22% ethanol
- Patulin levels in cider were reduced by fermentation but still exceeded 50 µg/kg, a maximum limit set by various regulatory agencies
- *Pa. niveus* was observed to be able to grow in low concentrations of three preservatives: potassium sorbate, sulfur dioxide, and sodium benzoate

## 1. Introduction

The mycotoxin patulin is an important apple-associated mycotoxin produced by a handful of fungi and noted for its genotoxic, immunosuppressant, and cytotoxic properties (Ülger et al., 2020). Patulin contamination in apple (Malus xdomestica) juices and ciders, subject to regulation in many parts of the world, is a long-standing issue that has been reported in various countries (Affairs, 2020; Commission, 2003; Commission Regulation, 2006; Harris et al., 2009; Spadaro et al., 2007; Yuan et al., 2010). It has been long assumed that patulin contamination of fruits and fruit products occurs through fruit infection by patulin-producing Penicillium species, particularly the notorious post-harvest pathogen *Penicillium expansum*.

While patulin has been detected in a variety of fruit products (Spadaro et al., 2007; Zouaoui et al., 2015), the mycotoxin is not traditionally thought to be an issue for their fermented counterparts. This is in part due to research that has shown that fermentation reduces patulin levels significantly (Erdoğan et al., 2018; Stinson et al., 1978; Zhang et al., 2019).

An alternative route of patulin entry into fruits and fruit products is through the notorious food-spoiling mold *Paecilomyces niveus*. This ascomycete produces heat-resistant ascospores that can survive lower temperatures of pasteurization and grow in acidic and low-oxygen conditions (Biango-Daniels et al., 2019; Biango-Daniels and Hodge, 2018; Taniwaki et al., 2009). A common soil fungus, *Pa. niveus* is particularly problematic for finished fruit products, and has been reported in apple puree, orange juice, and strawberry puree, raising concerns that it may be a source of patulin contamination in other juices like lemonade and orange juice (*Citrus* spp.) (Santos et al., 2018).

New research shows that *Pa. niveus* can grow as a post-harvest pathogen in various fruits, including apples (Biango-Daniels and Hodge, 2018; Wang and Hodge, 2022, 2020). Juice from infected fruits can become infested with living *Pa. niveus* mycelium and spores and contaminated with patulin (Biango-Daniels et al., 2019). Furthermore, this fungus can resist high temperatures, grow in low-oxygen environments, and resist fairly high concentrations of alcohol (Brown and Smith, 1957). The connection between patulin and post-fermented products is understudied (Al Riachy et al., 2021). But if *Pa. niveus* can survive and grow during fermentation, it may continuously produce patulin, contaminating the finished cider.

In this study, we investigate how each step of hard cider production impacts patulin-producing fungi introduced through infected fruit. We also evaluate the ability of common sanitation methods to decrease risks of spoilage and patulin contamination.

Cider apples are known for their higher tannin and phenol levels which have been hypothesized to be inhibitory to pathogens (Serrano et al., 2009). However, *Pa. niveus* has recently been found to be able to infect a variety of fruits beyond apples, so we hypothesize that cider apples too are susceptible to Paecilomyces rot (Wang and Hodge, 2022). Understanding how the different steps of hard cider production impact fungal growth and patulin production will help apple producers and hard cider makers better assess the risk of mycotoxin contamination.

Our objectives are to: 1. Test the susceptibility of traditional cider apple cultivars to four known patulin-producing strains of apple pathogens: *Penicillium expansum, Penicillium griseofulvum, Penicillium carneum*, and *Paecilomyces niveus*; 2. Assess the ability of each fungus to produce patulin in cider apple fruits; 3. Test the ability of *Pa. niveus* to produce patulin in lemonade and orange juice; 4. Test potential inhibitory effects of fermentation on the growth of the four fungi in small-scale bench-top fermentation by evaluating growth; 5. Develop and validate primers based on the RPBII gene for quantification of *Paecilomyces* sp. DNA; 6. Quantify the effects of three different preservative treatments and flash pasteurization on *Pa. niveus* growth and viability.

## 2. Materials and methods

To evaluate the impact of infected apples on patulin content of finished cider, we first prepared lab-infected apples of two traditional cider apple cultivars (Dabinett and Medaille d’Or) using four patulin-producing fungi. Two additional cider apple cultivars, Harry Masters Jersey and Chisel Jersey, were tested for *Paecilomyces niveus* susceptibility. We performed bench-scale cider fermentation, then evaluated two important attributes of the finished fermented cider: 1) patulin content, and 2) evidence of viable fungus. Any visible hyphae was grown and identified. To test the impact of preservatives on *Pa. niveus* spore germination and growth, ascospores underwent three preservative treatments in apple cider, each at three concentrations. Fungal growth of *Pa. niveus* was measured using qPCR, applying our newly designed PCR primers. *Paecilomyces niveus* spores in apple cider were additionally treated by capillary tube flash pasteurization, and spore viability was measured as colony forming units after plating on PDA.

### 2.1 Fungal cultures and apple infection

*Penicillium expansum* isolate 94222 was obtained from a department fungal collection. *Penicillium griseofulvum* NRRL 2159A and *Penicillium carneum* NRRL 25170 were obtained from the ARS NRRL fungal culture collection, and *Paecilomyces niveus* CO7 from the Hodge lab culture collection (Biango-Daniels et al., 2018). Fungi were grown in the dark at 25°C. To prepare toothpick inoculum, toothpick halves were tyndallized by autoclaving first in water, then in potato dextrose broth. They were placed on potato dextrose agar along with three to four plugs from the growing edges of each of the four fungal isolates. Control toothpicks were treated similarly but without the presence of fungi.

Once the toothpicks were fully colonized after two weeks, they were used to inoculate apples for cultivar trials and patulin experiments. Four cider apple cultivars were sourced from The Cornell Agricultural Experiment Station research orchard in Ithaca, NY(Chisel Jersey, Dabinett, Harry Masters Jersey, and Medaille d’Or) and inoculated following methods of Biango-Daniels and Hodge (2018). Briefly, apple fruits were first sanitized in 70% ethanol for 30 seconds, then two minutes in 1% sodium hypochlorite, and finally in 70% ethanol for 15 seconds. Fruits were numbered and left to dry in a class IIB biological safety cabinet. Infested toothpicks were inserted 1 cm deep into the fruit.

### 2.2 Bench-top fermentation

Toothpicks used in inoculation were removed from roughly 4.5 kg of Dabinett and Medaille d’Or cider apples infected 4 weeks prior with *Pe. expansum*. The apples were chopped and homogenized with a blender, and the puree was strained through 4 layers of cheesecloth. Samples of 150mL of cider were aliquoted in three separate 500mL flasks. The remaining cider was frozen for brix and patulin quantification. One mL (1.13g of yeast and emulsifier) of activated *Saccharomyces cerevisiae* strain EC1118 (Lalvin) prepared under manufacturer’s instructions was added to each of the three flasks. This process was repeated for fruits infected with *Pe. griseofulvum, Pe. carneum*, and *Pa. niveus*. Flasks were sealed with autoclaved rubber bungs that were fitted with air locks. The flasks of cider were left to ferment at room temperature for four weeks and CO2 bubbling was monitored daily. Preservative treatments were not used during fermentation but were tested later. At four weeks, fermented product was checked for fungal growth and any present hyphae was grown and identified. As a baseline, uninfected Dabinett and Medaille d’Or apples stored at 25°C in the dark for four weeks were processed and fermented in the same way.

### 2.3 Patulin quantification

Lemonade, orange juice, and apple cider used in the pasteurization and preservative treatment protocols were free from preservatives and purchased from a local supermarket in New York. For patulin analysis of infested lemonade, orange juice and apple cider, 50mL aliquots were submitted to Trilogy Analytical Laboratory (Washington, MO). Patulin was quantified by high-performance liquid chromatography following official AOAC International methods for apple juice (Brause et al., 1996).

### 2.4 Impact of common preservatives on Paecilomyces niveus spore germination and growth

*Paecilomyces niveus* ascospore viability was tested using three food-relevant treatment conditions designed to limit microbial growth: potassium sorbate, sulfur dioxide, and sodium benzoate. Food-grade apple cider was aliquoted into 2-mL tubes infested with 2 × 105 *Pa. niveus* spores harvested from 4-week-old cultures on PDA. For each of the three preservatives, three different concentrations were tested. Tubes were then capped and air sealed. Acidity of the cider was measured with a digital pH meter and found to be 3.6. Food grade potassium sorbate was weighed to create three concentrations of potassium sorbate at .02%, .06%, and .1%. Campden tablets were crushed and weighted to create .05%, .1% and .15% treatments of sulfur dioxide in apple cider. Food grade sodium benzoate was weighed to create .05%, .1%, and .15% concentrations. Positive controls lacked any preservative treatment, and negative controls had neither spore treatment nor preservative treatment. Each treatment was performed in triplicate and kept in the dark at 25°C. After 2 weeks, presence or absence of hyphal growth in each sample was visually assessed. Present hyphae was extracted, grown on PDA, and identified morphologically. Samples were then centrifuged at 16,000 xg for 5 minutes and DNA extraction was performed on the resulting pellet.

### 2.5 RPBII primer design and qPCR validation

Primers and a qPCR system were designed for accurate quantification of *Paecilomyces niveus* DNA after exposure to various treatments. Low-coverage genomes of 23 *Pa. niveus* isolates were visualized using NCBI Genome Workbench and compared to the high-coverage genome of *Pa. niveus* isolate CO7 (Biango-Daniels et al., 2018). Primers were designed using the PrimerQuest Tool (https://www.idtdna.com/primerquest/Home/Index) and candidate sequences within the RNA polymerase II gene were compared to sequences of *Pa. fulvus* and *Pa. variotii* aligned using MUSCLE (Edgar, 2004) and visualized using Snapgene software (https://snapgene.com/). Primer specificity was additionally verified through BLAST. Primers were tested using PCR on DNA extracts of 9 food-spoiling and post-harvest fruit pathogens, 3 isolates each of *Pa. fulvus* and *Pa. variotii*, and 24 isolates of *Pa. niveus*. Fungal cultures were obtained from Hodge lab collections at Cornell University and from the NRRL Culture Collection.

Genomic DNA was extracted from fungal isolates using the DNeasy PowerSoil Pro Kit (Qiagen) following manufacturer instructions. Q5 High Fidelity DNA Polymerase (New England Biolabs, United States) was used for end point PCR to determine specificity of the PaeRPB2f/r primers (Table 2) under the following PCR cycles: 98°C for 30s and 26 cycles of 98°C for 10s (denaturation), 60°C for 30s (annealing), and 72°C for 45s (extension). For *Pa. niveus* DNA quantification, the CFX Connect Real-Time PCR Detection system (Bio-Rad, Hercules, CA, USA) was programmed to run at 95°C for 10 minutes and then to complete 40 cycles of 95°C for 15s (denaturation), 60°C for 30s (annealing), and 72°C for 45s (extension). All reactions were done in triplicate and results were measured by quantification cycle (Cq).

**Table 1.**
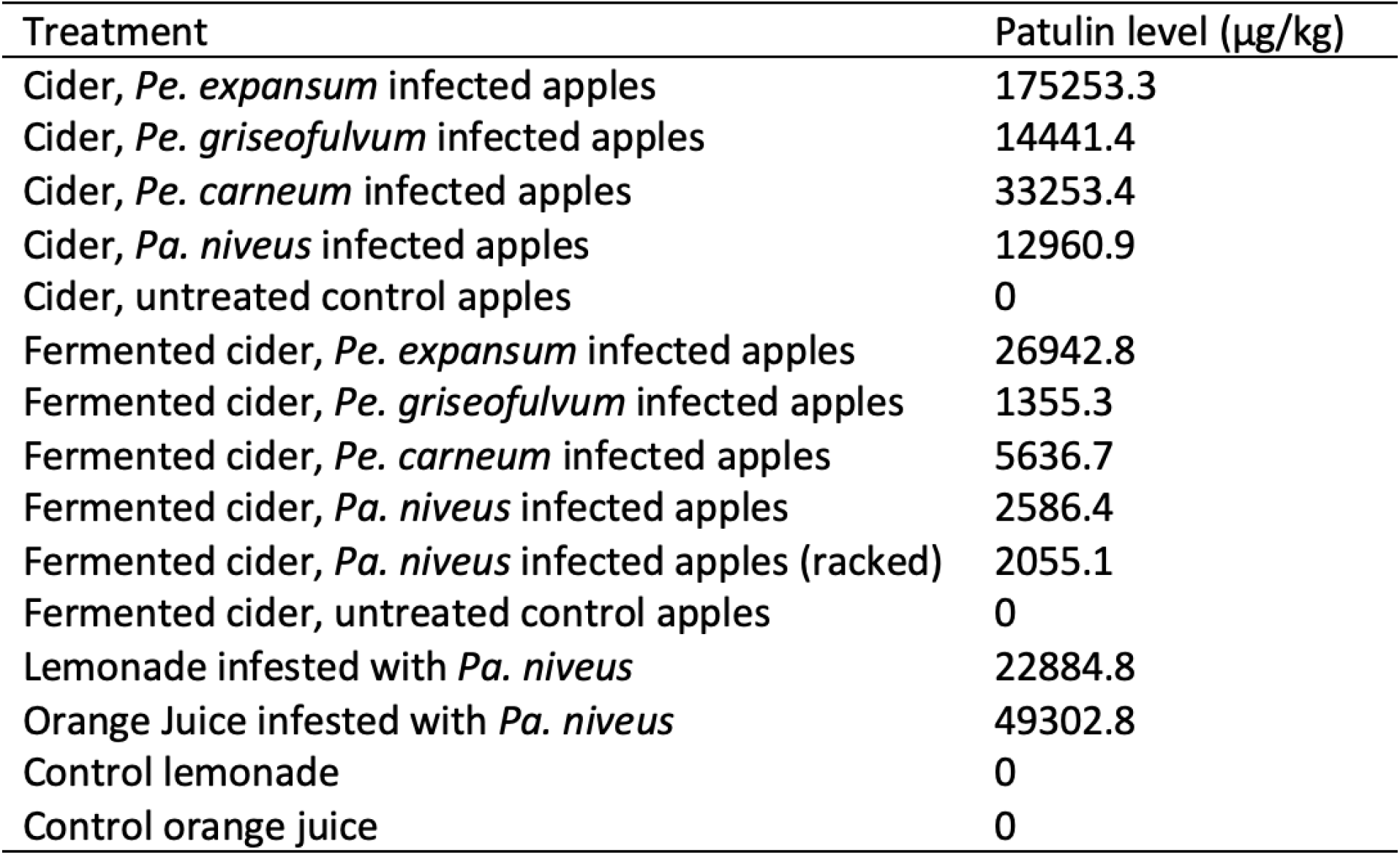
Patulin concentrations (µg/kg) in cider and juices. Patulin was quantified from cider extracted from infected cider apples both before fermentation (n=1) and after fermentation (n=1). Yeast cells were removed (racked) by decanting clear cider from the sediment of the *Pa. niveus* infected cider apple sample. Both racked and unracked samples were quantified for patulin. 200 mL store-bought lemonade and orange juice were infested with 1x 10^7 *Pa. niveus* asci/ascospores and patulin was quantified (n=1) after two weeks. The limit for patulin contamination in the United States and Europe for fruit juices is 50 µg/kg (Affairs, 2020; Commission, 2003; Commission Regulation, 2006).

| Treatment | Patulin level (µg/kg) |
| --- | --- |
| Cider, *Pe. expansum* infected apples | 175253.3 |
| Cider, *Pe. griseofulvum* infected apples | 14441.4 |
| Cider, *Pe. carneum* infected apples | 33253.4 |
| Cider, *Pa. niveus* infected apples | 12960.9 |
| Cider, untreated control apples | 0 |
| Fermented cider, *Pe. expansum* infected apples | 26942.8 |
| Fermented cider, *Pe. griseofulvum* infected apples | 1355.3 |
| Fermented cider, *Pe. carneum* infected apples | 5636.7 |
| Fermented cider, *Pa. niveus* infected apples | 2586.4 |
| Fermented cider, *Pa. niveus* infected apples (racked) | 2055.1 |
| Fermented cider, untreated control apples | 0 |
| Lemonade infested with *Pa. niveus* | 22884.8 |
| Orange Juice infested with *Pa. niveus* | 49302.8 |
| Control lemonade | 0 |
| Control orange juice | 0 |

**Table 2.**
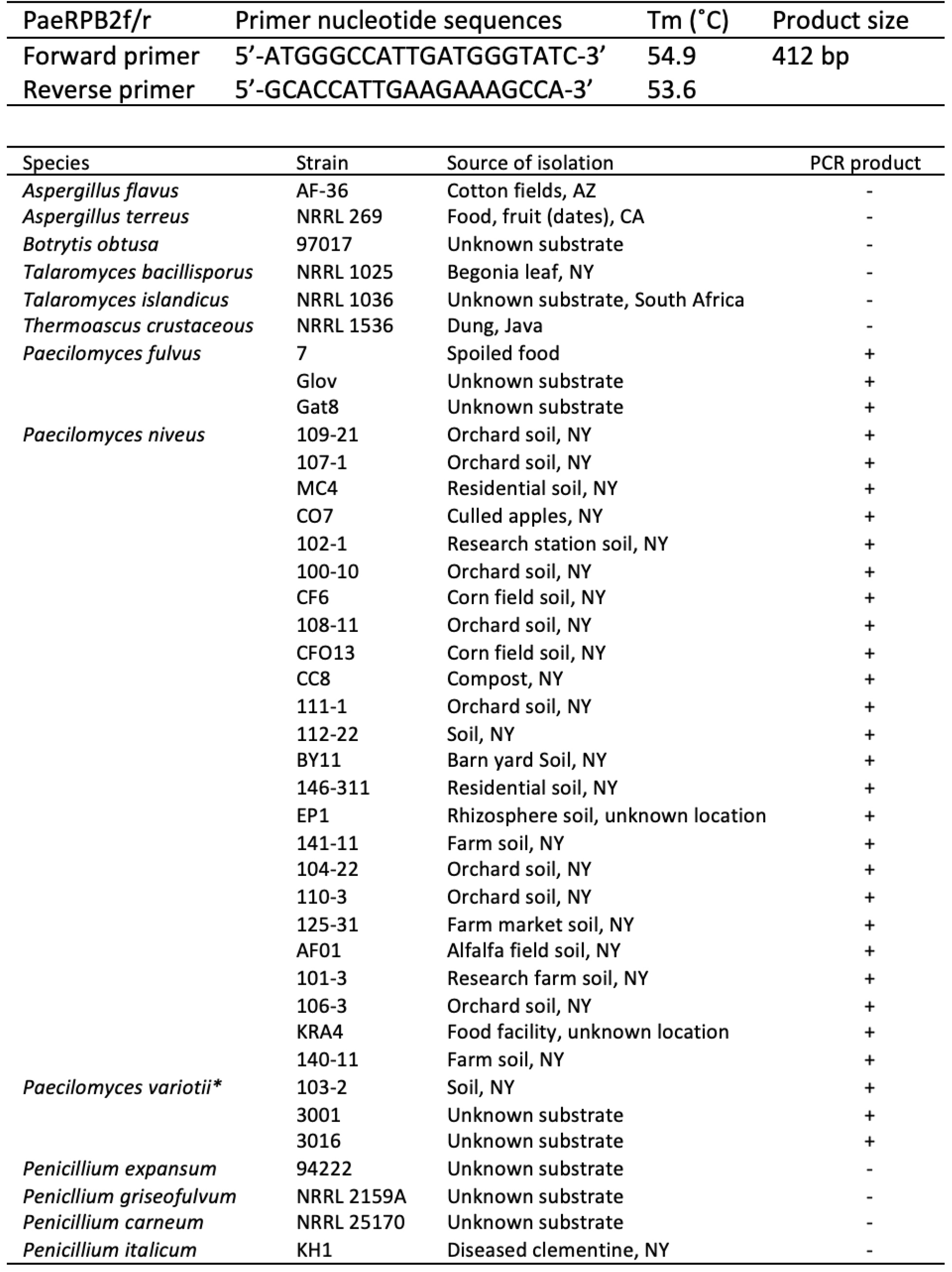
Sequences and specificity of RPBII primers designed in this study for quantification of *Pa. niveus* growth using real-time PCR. End-point PCR was performed to test the specificity of the primers to *Paecilomyces* species.

qPCR reaction mixtures contained 25µl 2x Power SYBR Green PCR Master Mix (Applied Biosystems, Madrid, Spain), 1µl 10µM PaeRPB2f primer, 1µl 10µM PaeRPB2r primer, 10µl of eluted fungal DNA, and 13 µl deionized water (50µl total). Real time qPCR was performed on the CFX Connect Real-Time PCR System (Bio-Rad, Hercules, CA, USA) programmed to hold at 95°C for 10 minutes followed by 40 cycles of 95°C for 15s (denaturation), 60°C for 30s (annealing), and 72°C for 45s (extension).

### 2.6 Heat-treatment

*Paecilomyces niveus* asci and ascospores were extracted from 4-week-old plates on PDA using water and autoclaved pipette tips to agitate the surface of the mycelia. The spore solution was filtered through 8 layers of cheesecloth and one of Whatman No. 1 filter paper to remove hyphae. A haemocytometer was used to determine spore concentration and to confirm absence of hyphal fragments. Apple cider free from preservatives was purchased from a local supermarket, autoclaved and spiked with *Pa. niveus* asci and ascospores. 20µl samples containing roughly 200 *Pa. niveus* colony forming units were aliquoted into 75mm long thin walled (.2 ± .02 mm thick) soda glass capillary tubes (Kimble Chase). The empty ends of the capillary tubes were sealed using fire. Each was exposed to either a light heat-treatment of 71.1°C for 6 seconds (n=14), a heavy treatment of 71.7°C for 15 seconds (n=14) or no treatment (n=14). The light treatment is based on the United States Food and Drug Administration (FDA) minimum pasteurization process for non-shelf stable juices, and the heavy treatment is based on the US protocol for flash pasteurization of milk, both of which are used for log reduction of harmful pathogens. After heat-treatment, the capillary tubes were cooled in an ice bath for 3 seconds, broken, and spread plated on Rose Bengal Agar. Colony formation was observed and quantified over the next two weeks.

### 2.7 Statistical analysis

Significance of cider apple lesion development was tested by comparing lesion sizes via two separate linear mixed-effects models at day 2 and day 8 for both apple cultivars infected by *Pe. griseofulvum* and *Pa. niveus* (cider apples infected with *Pe. expansum* and *Pe. carneum* were tested at days 3 and 8) and by comparing lesion sizes from both the control and treatments on day 8. Apple identification number was accounted for as a fixed effect. Variation in pathogenicity by *Pa. niveus* infection in apple cultivars was evaluated by one-way analysis of variance using diameter of lesions on day 20. Differences between heat-treatments on *Pa. niveus* spore viability and colony forming units were tested using the Kruskal-Wallis rank sum test. Statistical analysis was followed by a post-hoc Tukey Honest Significant Difference Test. Statistical models were executed in R statistical programming software using the emmeans, lme4, lmerTest, and stats packages. Data was visualized using the ggpubr and ggplot2 packages.

## 3. Results

### 3.1 Characterization of Paecilomyces niveus and three Penicillium sp. infections on cider apples

In cider apples infected by *Pe. expansum*, dark red-brown circular lesions in Dabinett (Fig. 1B) or brown lesions in Medaille d’Or apples (Fig. 1C) rapidly developed at the site of the inoculation. Apple flesh quickly lost integrity as soft rot developed, particularly with infected Dabinett apples. In Medaille d’Or apples, concentrated tufts of sporulating hyphae appeared on the lesion surface by 20 days post-inoculation.

**Figure 1.**
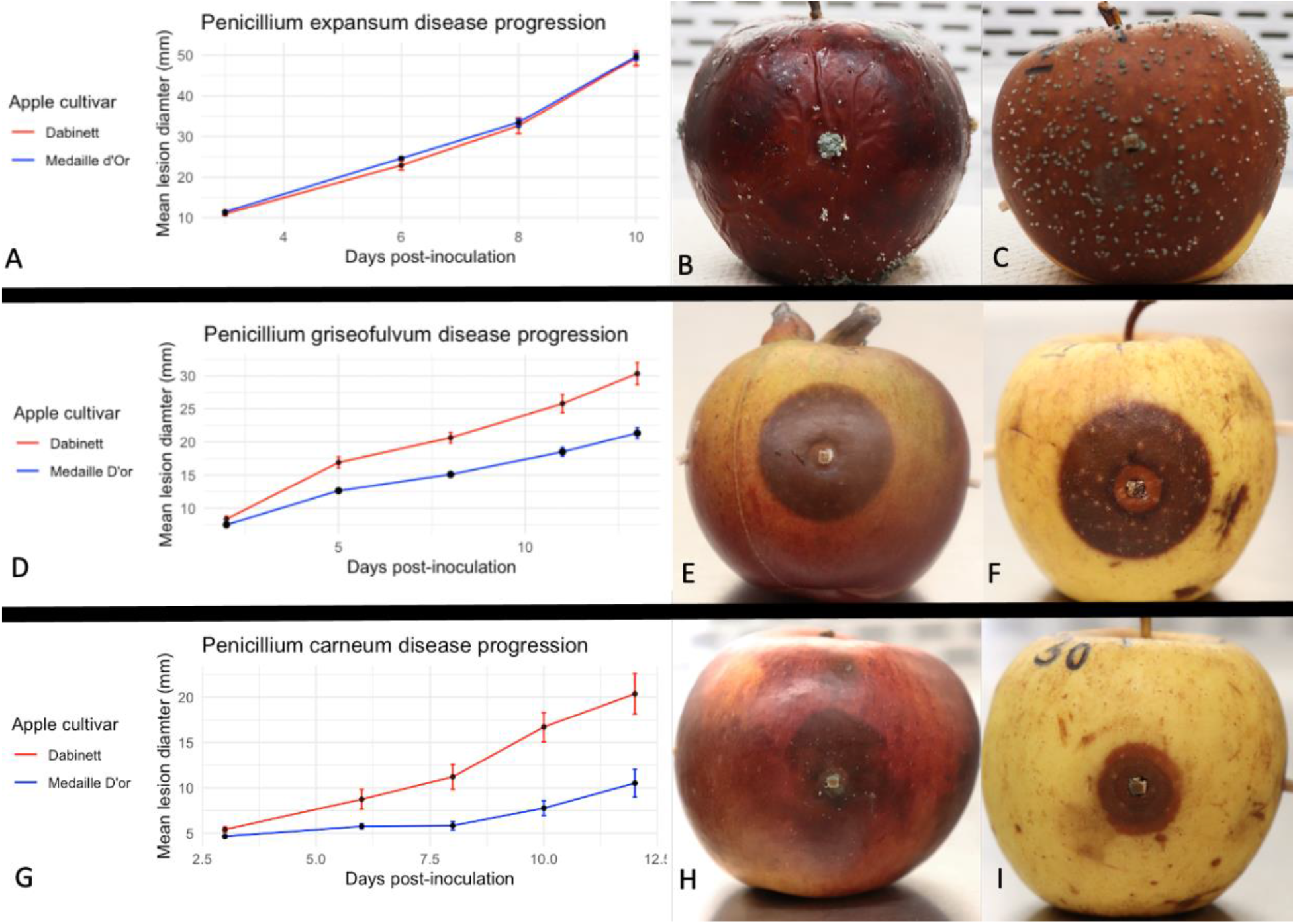
Disease progress and external symptoms of two cider apple cultivars 3 weeks after inoculation with three *Penicillium* spp. and incubation in dark, moist chambers (25°C, ≥95% humidity). Top row shows *Penicillium expansum* lesion diameters (A) and infections of Dabinett (B) and Medaille d’Or (C). Middle row shows *Penicillium griseofulvum* lesion diameters (D) and infections of Dabinett (E) and Medaille d’Or (F). Bottom row shows *Penicillium carneum* lesion diameters (G) and infections of Dabinett (H) and Medaille d’Or (I).

Both cultivars (Fig. 1E, 1F) of cider apples infected with *Pe. griseofulvum* developed expanding brown lesions centered at the point of inoculation that were sometimes lighter brown close to the point of inoculation. Lesion borders were distinct and pronounced in both cultivars. Rot within the apple is light brown and unlike the soft rot infections by *Penicillium expansum*, apple flesh infected with *Pe. griseofulvum* was firm and spongy to the touch.

Cider apples infected with *Pe. carneum* developed brown to light brown lesions and soft rot in both Dabinett and Medaille d’Or cultivars (Fig. 1H, 1I). Lesion borders were often diffuse in Dabinett apples. Lesions sometimes displayed concentric circles with varying shades of brown in Medaille d’Or apples. At 20 days post-inoculation, a faint blue ring of spores was seen developing at the border of the endocarp within infected Dabinett apples.

All four cider apple cultivars (Chisel Jersey, Dabinett, Harry Masters Jersey, and Medaille d’Or) tested for susceptibility to *Pa. niveus* infection developed circular dark brown lesions at the point of inoculation (Fig. 2A). Lesion borders were generally sharp and distinct in Dabinett and Chisel Jersey apple fruits. Chlorosis sometimes developed near the edges of the lesions in Medaille d’Or fruits. Brown concentric circles sometimes developed, most clearly in Medaille d’Or and Harry Masters Jersey apples. Apple rot in all cultivars was firm to the touch and resulted in brown to light-brown internal rot.

**Figure 2.**
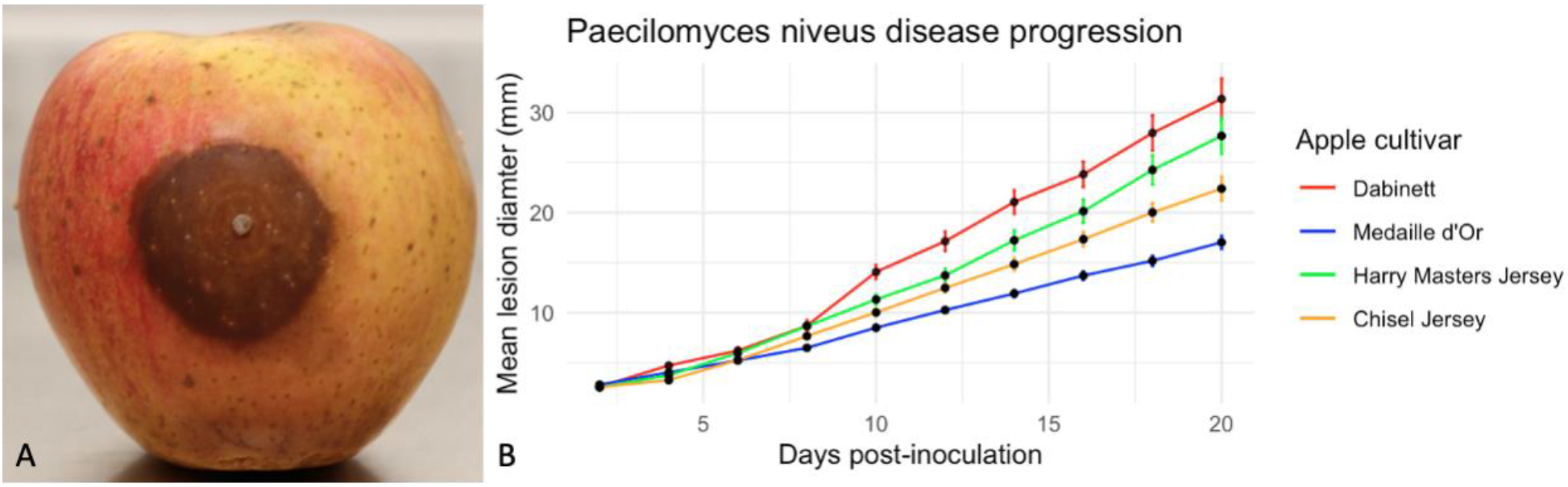
Paecilomyces rot in four cider apple cultivars. A, Harry Masters Jersey apple infected by *Pa. niveus*, 3-weeks post-inoculation. B, Line plot shows increase in mean lesion diameter over 3 weeks in four cider apple cultivars.

Our results show that each cider apple cultivar tested was susceptible to each of the four patulin-producing fungi, and that apple fruits responded differently to each infection. Infections by *Pe. expansum* and *Pe. carneum* elicited brown lesions and soft rot, symptoms consistent with Blue Mold infections. Infection by *Pe. griseofulvum* was unusual as it resulted in rot with a spongy texture, not a soft rot. This may be a point of interest for apple producers who may look specifically for Blue Mold infections in apple fruits. Additionally, because we used *Pe. griseofulvum* isolate NRRL 2159A, a white mutant, we did not see blue sporulation as a sign of the fungus. In *Pa. niveus* infections, symptoms of hard rot with occasional concentric circles and slowly spreading lesions were consistent with Paecilomyces rot symptoms in dessert apples (Biango-Daniels and Hodge, 2018). Follow-up studies may consider testing additional isolates of *Pa. niveus* and each *Penicillium* species for infection assays.

### 3.2 Susceptibility to patulin-producing apple pathogens

Inoculations by *Pa. niveus* and all three *Penicillium* species (*Pe. expansum, Pe. griseofulvum*, and *Pe. carneum*) resulted in clear lesion development in each cider apple cultivar tested within 3 to 6 days post-inoculation. The average of horizontal and vertical lesion diameters of each apple fruit’s two lesions was measured every 2 to 3 days. Lesion diameters (± standard error) grew rapidly over the course of the next 10 to 20 days. Between 3 and 8 days post-inoculation in apples infected with *Penicillium expansum*, average lesion diameter increased by 21.66 ± 1.95 mm (*n* = 21) in Dabinett apples, and 22 ± .83 mm (*n* = 29) in Medaille d’Or apples (Fig. 1A). In apples infected with *Pe. griseofulvum*, average lesion diameter increased by 12.06 ± .91 mm (*n* = 22) and 7.61 ± .48 mm (*n* = 28) in Dabinett and Medaille d’Or apples, respectively, between 2 and 8 days post-inoculation (Fig. 1D). Average lesion diameters in apples infected with *Pe. carneum* increased by 5.81 ± 1.41 mm (*n* = 22) and 1.17 ± .50 mm (*n* = 30) in Dabinett and Medaille d’Or apples, respectively, between 3 and 8 days post-inoculation (Fig. 1I). The average change in lesion diameter for cider apples infected with *Pa. niveus* (± standard error) was 5.09 ± .28 mm (*n* = 27) in Chisel Jersey apples; 3.67 ± .16 mm (*n* = 38) in Medaille d’Or apples; 6.12 ± .23 mm (*n* = 27) in Harry Masters Jersey apples; and 6.09 ± .57 mm (*n* = 40) in Dabinett apples between 2 and 8 days post-inoculation (Fig. 1G). Apples observed to be colonized by other fungi were removed from the experiment and from statistical analysis. On day 8, lesion diameters were significantly larger than both control lesions (*P* < .01), and treatment lesions on day 2 (*Pe. griseofulvum* and *Pa. niveus*) and day 3 (*Pe. expansum* and *Pe. carneum*) (*P* < .01). In addition, by one-way analysis of variance of lesion diameters of *Pa. niveus*-infected apples on day 20, apple cultivar was found to be significant (*P* < .01).

Neither Dabinett nor Medaille d’Or cider apple cultivars have been previously evaluated for susceptibility to *Pa. niveus* infection, nor to Blue Mold infection by *Pe. expansum, Pe. griseofulvum*, and *Pe. carneum*. Despite cider apple cultivars being known for their high tannin and phenol content, which have been proposed to confer some resistance to fungal infection (Serrano et al., 2009), all four cultivars were susceptible to infections by the four patulin-producing fungi. Our data suggest there may be some cultivar-based resistance to infections by *Pa. niveus* and raise the possibility of using more resistant cultivars as a strategy to combat patulin contamination arising from Paecilomyces rot. Previous research has demonstrated that patulin functions as a pathogenicity agent in *Pe. expansum*, facilitating partial lesion formation and breaking down fruit tissue (Bartholomew et al., 2022; Snini et al., 2016). While it is unclear whether patulin functions similarly for other postharvest pathogens, future studies may consider expanding on the scope of this work by exploring additional cultivars for resistance to Blue Mold and Paecilomyces rot and comparing pathogenicity of patulin producing and non-patulin producing isolates.

### 3.3 Koch’s postulates

For each of the four fungal pathogens, Koch’s postulates were satisfied using four-week post-inoculation Medaille d’Or and Dabinett apples. Diseased apple surfaces were sanitized with 70% ethanol, and diseased interior tissue extracted from apple lesions was plated on PDA. Fungal isolates were confirmed morphologically and through Sanger sequencing, and an isolate of each species was used to reinfect healthy apples. The same symptoms were observed, thereby satisfying Koch’s postulates and confirming the causal agent of the observed disease.

### 3.4 Benchtop fermentation and patulin quantification

After four weeks of fermentation, no mycelial growth was observed in the control flasks, nor in fermented cider extracted from apples infected with *Pe. expansum, Pe. griseofulvum*, or *Pe. carneum*. However, mycelial growth was present in all three replicates of cider extracted from apples infected with *Pa. niveus*. Mycelia was extracted and grown on PDA and confirmed morphologically to be *Pa. niveus*. Soluble solid concentration values were 10.13, 9.77, 10.63, 11.67, and 10.6 °Brix before fermentation and 2.9, 3.5, 4, 3.4, and 4.5 °Brix after four weeks of fermentation for cider samples made from uninfected apples and apples infected with *Pe. expansum, Pe. griseofulvum, Pe. carneum, Pa. niveus* respectively.

After four weeks of fermentation, cider made from apples infected with *Pe. expansum* exhibited the highest patulin concentration at 175253.3 µg/kg, while cider from apples infected with *Pa. niveus* had the lowest concentration of patulin at 12960.9 µg/kg (Table 1). We observed post-fermentation reduction in patulin levels in all four cider samples extracted from infected apples. The largest reduction in patulin was observed in cider extracted from apples infected with *Pe. griseofulvum* (90.6%) while the smallest reduction occurred with cider extracted from apples infected with *Pa. niveus* (80.0%). In all cider samples before and after fermentation, patulin levels far exceeded the United States FDA and Europe limit of 50 µg/kg (Affairs, 2020; Commission, 2003; Commission Regulation, 2006).

Store bought lemonade and orange juice, in 200 mL aliquots, were each spiked with *Pa. niveus* 10^7 asci and ascospores and left to sit at room temperature. A surface layer of white mycelium was observed developing in the treated samples. Patulin was quantified after two weeks at 22884.8 µg/kg and 49302.8 µg/kg for lemonade and orange juice respectively.

In this study, we observed that fermentation does not completely inhibit the growth of *Pa. niveus* in cider, which raises the possibility of spoilage of finished hard ciders by the fungus. Patulin reduction after fermentation was smallest in cider samples infested with *Pa. niveus*. Our results suggest this may be due to growth and production of patulin by *Pa. niveus*. Our data suggest *Pa. niveus* is also able to produce patulin in lemonade and orange juice. Although our study did not include a control for the confounding variable of natural patulin loss in the absence of yeast, previous work has shown this form of reduction is minimal (Stinson et al., 1978).

### 3.5 RPBII primer design for detection of Paecilomyces spp

Primers based on the RNA polymerase II gene were designed and tested on multiple strains of three different *Paecilomyces* species: *Pa. niveus* (n=24), *Pa. fulvus* (n=3), and *Pa. variotii* (n=3), in addition to nine close relatives, including postharvest pathogens and food spoiling molds in the order Eurotiales (Table 2). PCR products were obtained from each of the three *Paecilomyces* species, but not from any of the other Eurotiales fungi tested.

### 3.6 Effects of industrial preservative treatments on Paecilomyces niveus spore survival and growth

To determine the effects of industry standard food treatment on *Paecilomyces niveus* spore survival and growth, food-grade apple cider infested with *Pa. niveus* CO7 spores was subjected to treatments commonly used to control human disease and food spoilage agents. Treatments of potassium sorbate, sodium benzoate, and sulfur dioxide were applied in three concentrations, and fungal growth was assessed visually for presence of hyphae. As a proxy for *Pa. niveus* biomass, DNA was quantified by qPCR using primers developed in this study.

After two weeks of exposure, *Pa. niveus* hyphal growth was observed in 2mL tubes containing food-grade apple cider, as well as in low concentrations of each of the three preservatives (.02% potassium sorbate, .05% sodium benzoate, and .05% sulfur dioxide). Hyphae were also observed at .1% concentration of sulfur dioxide, but growth was not detected at higher concentrations of the three preservative treatments (Table 3). Mean Cq values (n=3) estimating *Pa. niveus* DNA in the 2mL tubes were obtained for each treatment. Higher Cq values were generally observed for tubes treated with high preservative concentrations, suggesting less *Pa. niveus* biomass (Table 3).

**Table 3.** *Pa. niveus* germination and growth in apple cider treated with low amounts of potassium sorbate, sodium benzoate, and sulfur dioxide, as observed by visible hyphal growth and mean Cq values, a proxy for fungal biomass. Mean Cq and standard deviation values from real-time qPCR assays on 2mL aliquots of apple cider infested with *Pa. niveus* spores after a 2-week exposure to industry standard preservative treatments.

| Treatment | Concentration (%) | Observed Hyphal Growth | Mean Cq values |
| --- | --- | --- | --- |
| Cider negative control | N/A | - | N/A |
| Cider positive control | N/A | + | 22.59 ± .82 |
| Potassium sorbate (Low) | .02 | + | 21.53 ± 1.94 |
| Potassium sorbate (Medium) | .06 | - | 28.83 ± 4.50 |
| Potassium sorbate (High) | .1 | - | 30.87 ± 5.99 |
| Sodium benzoate (Low) | .05 | + | 22.33 ± 1.77 |
| Sodium benzoate (Medium) | .1 | - | 25.48 ± 1.11 |
| Sodium benzoate (High) | .15 | - | 26.80 ± 1.41 |
| Sulfur dioxide (Low) | .05 | + | 24.40 ± 3.86 |
| Sulfur dioxide (Medium) | .1 | + | 20.94 ± 1.37 |
| Sulfur dioxide (High) | .15 | - | 25.64 ± 1.93 |

*Paecilomyces niveus* spores that underwent two flash pasteurization protocols in soda glass capillary tubes were assessed for colony forming units. In both treatments, a significant number of *Pa. niveus* spores remained viable (Figure 3). There was no significant difference between the 71.1°C 6 second treatment and the 71.7°C 15 second treatment under the Kruskal-Wallis rank sum test (χ2 = 5.405, df = 2, p=.067).

**Figure 3.**
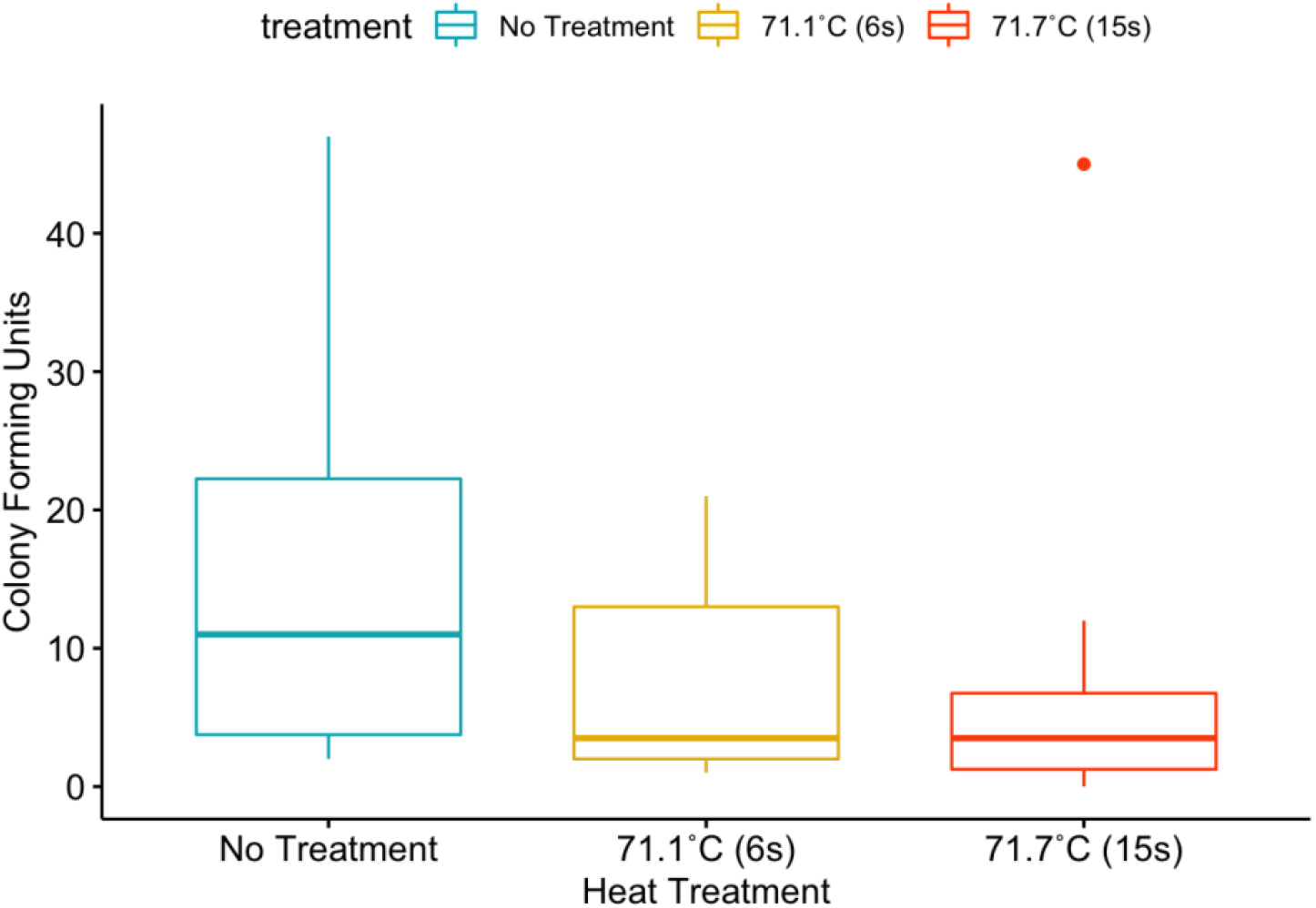
Box plots showing the impact of heat treatment on spore viability. The number of colony-forming units is shown for three treatments. Roughly 200 asci/ascospores in 20µl glass capillary tube aliquots of food-grade apple cider were exposed to 6s at 71.1°C, 15s at 71.7°C (n=14), or no treatment.

## 4. Conclusions and Discussion

This study is the first to report on the susceptibility of cider apples to Paecilomyces rot. In addition, we also report on the susceptibility of two cider apple cultivars Dabinett and Medaille d’Or to *Penicillium expansum, Penicillium griseofulvum*, and *Penicillium carneum* infections. Cider apple cultivars typically contain higher levels of polyphenolic compounds, which are known to have antimicrobial ability, yet each of the four fungi were able to infect and grow in all cider cultivars tested (Marks et al., 2007; Serrano et al., 2009). These results are consistent with previous pathogenicity assays of *Pe. expansum* with traditional apple cultivars (Lončarić et al., 2021). Patulin has been previously reported as a cultivar-dependent virulence agent for *Pe. expansum* infections of apples and future studies could investigate if patulin serves a similar role in infection for other patulin-producing pathogens (Snini et al., 2016).

Hard cider makes up a significant portion of the beverages in several countries and is a growing industry in the United States (Ewing and Rasco, 2018). Traditional cider production and apple harvesting, especially in European countries, may employ machinery to shake trees before sweeping up “drop” fruits that fall to the ground, a practice referred to as “shake and sweep” (Karl et al., 2022; Miles et al., 2020). As drop fruits may be both wounded and exposed to soil, they may introduce both foodborne disease organisms and spoilage inoculum (Ewing and Rasco, 2018; Guo et al., 2002), including the patulin-producing fungi highlighted here. Future research could investigate risk factors for introducing patulin-producing spoilage inoculum in hard ciders that originate from practices related to fruit harvesting and processing.

Our results demonstrate that *Pa. niveus*, unlike the other three tested *Penicillium spp*., can not only survive apple processing and bench-top fermentation, but can also grow in low-oxygen, finished hard cider product. Food spoilage by *Pa. niveus* is a long-standing problem for fruit juices and products and has been found in apple juice production facilities and in various fruit products (Salomão et al., 2014; Santos et al., 2018). *Paecilomyces niveus* ascospores can also survive high temperatures, making this food-spoiling agent troublesome for fruit processing facilities (Biango-Daniels et al., 2019; Taniwaki et al., 2009).

Previous studies have explored strategies for patulin mitigation in solid foods and fruit juices (Ioi et al., 2017; Moake et al., 2005). Alcoholic fruit products are however not traditionally considered high-risk for patulin contamination, in part because fermentation has been shown to significantly reduce patulin levels (Erdoğan et al., 2018; Stinson et al., 1978; Zhang et al., 2019). Our results raise the possibility that *Pa. niveus* spoilage inoculum remains a human health hazard in hard ciders, in that it may survive fermentation and grow in the finished fermented product. This concern extends to other fruit juice products as we observed patulin levels far above the 50 µg/kg in lemonade and orange juice inoculated with *Pa. niveus* spores. To help quantify the risk of *Pa. niveus* spoilage in hard cider products, next steps should include surveying patulin contamination and the incidence of *Pa. niveus* spoilage in finished ciders.

Bioassays testing the effectiveness of sulfur dioxide, potassium sorbate, and sodium benzoate showed variable effectiveness at restricting on *Pa. niveus* growth after two weeks. We observed that under low (.05%) and moderate (.1%) concentrations of sulfur dioxide, significant *Pa. niveus* growth was noted as the presence of fungal hyphae. However, while sulfur dioxide is commonly used as preservatives to control fungal and bacterial growth in fruit juices, their effectiveness is highly pH dependent due to the mode of action of sulfurous acid (Silva and Lidon, 2016). It is possible that the pH of the apple cider we used (pH = 3.6) would need to be lowered for effective sulfur dioxide treatment. Both sodium benzoate and potassium sorbate, the salts of organic acids (benzoic acid and sorbic acid respectively), are also extensively used as preservatives in low pH foods (Silva and Lidon, 2016), and potassium sorbate is additionally used to help stop post-fermentation of unfermented sugars, especially in wines (Lück, 1990; Silva and Lidon, 2016). Moderate levels of both organic acids (.1% sodium benzoate and .06% potassium sorbate) in food-grade apple cider were observed to effectively restrict *Pa. niveus* hyphal growth over two weeks, compared to the control.

Results from this study help assess potential patulin contamination of apple cider and hard cider by four patulin-producing apple pathogens, especially *Pa. niveus*. Our work also introduces a primer pair specific to the RPBII region of *Paecilomyces spp*. and a qPCR protocol to detect and quantify three troublesome food spoilage fungi: *Pa. niveus, Pa. fulvus*, and *Pa. variotii*. qPCR methods introduced here can be applied to future bioassays involving these three *Paecilomyces spp*. Patulin is a regulated mycotoxin by various regulatory agencies and risk of contamination is both an issue for consumer health and food production concern (Affairs, 2020; Commission, 2003; Commission Regulation, 2006). These insights raise a concerning potential connection between patulin and hard cider and will aid in optimizing apple fruit and cider production with respect to food safety and consumer health.

## Acknowledgements

We gratefully acknowledge the NRRL ARS culture collection for providing multiple fungi used in this paper. We also thank the Melanie J. Filiatrault lab for sharing expertise on qPCR, the Randy W. Worobo Lab for their assistance in obtaining cultures of *Paecilomyces* spp. and Aarzoo Haideri and Maryann McCloskey for assistance with bioassays and heat treatment protocols.

